# *ALGR*: A multi-purpose agricultural landscape generator in R

**DOI:** 10.1101/2025.03.31.646291

**Authors:** Eyal Goldstein, Antonia Deutscher, Eamon O’Keeffe, Kerstin Wiegand

## Abstract

Agricultural and ecological modelers commonly use maps as input for spatially explicit simulations. While real world maps are often used, they are limited by being static objects, therefore making it difficult to assess how patterns within the landscape contribute to ecological processes. Agricultural landscape generators (ALG) are a useful tool for simulating maps in a more flexible way. They can increase robustness of models that rely on landscape maps as input, they allow modelers to give spatial representation to non-spatial models, and they are a useful tool for recreating spatial patterns in agricultural-dominated landscapes. A limitation of previous *ALGs* is that they have rarely been designed for general use (non-open source software, not written in R, and designed for specific projects). Furthermore, they are typically either extremely general and thus oversimplified or have a high specificity for particular use cases.
*ALGR* bridges this gap by providing a general-purpose, dynamic landscape generator that balances structural realism with adaptability. *ALGR* generates agricultural landscapes with a three-step approach: first, outlining potential space, second, field placement inside of that space, and third, enrichment of the landscape with information. This stepwise approach ensures that *ALGR* generates landscapes with realistic spatial patterns while remaining adaptable to diverse regions and applications.
It is the first ALG that is specifically designed to allow a simple integration within the R programming environment and the *r-spatial* package environment. *ALGR* is designed as a general-purpose generator, which is simple to use and facilitates an easy integration in modelling workflow.
We present several examples of workflows using *ALGR*, to demonstrate its usefulness. Our examples include: 1) simulating different land use shares, 2) parameter tuning of *ALGR* to recreate real world landscape patterns 3) spatially distributing crop portfolios, and 4) using real world maps as a basis for field placement

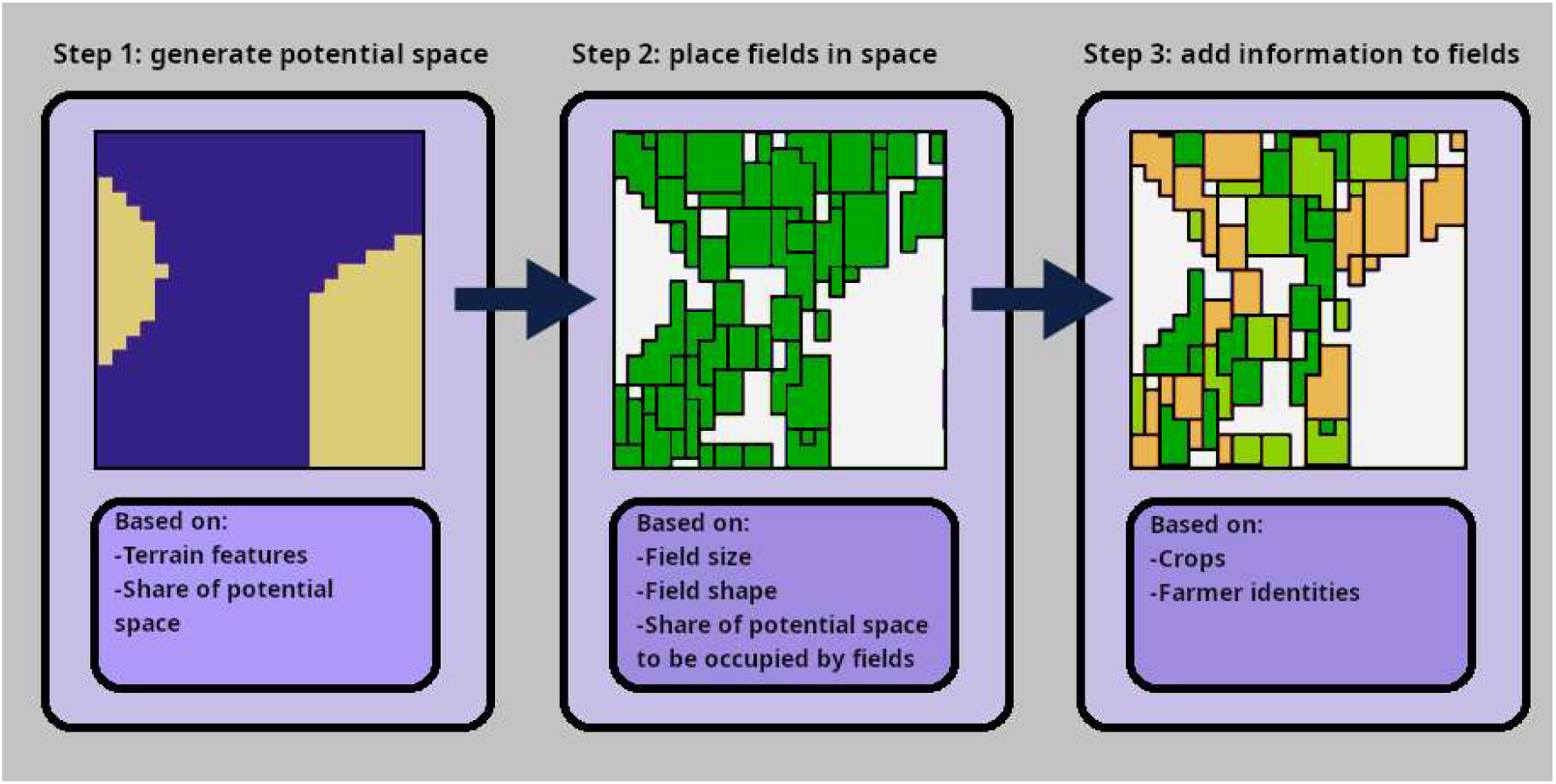

Visual Abstract: The three-step conceptual framework of *ALGR* for landscape generation

## Introduction

Landscape maps are fundamental to agricultural and ecological modeling, providing the spatial structure necessary for simulating ecological and economic processes. Use of landscape maps can be found in agent-based models (Becher et al., 2014; Iwamura et al., 2014; Lambin et al., 2010), mechanistic models (Häussler et al., 2017; Martín et al., 2024; Poggi et al., 2021), as well as different bio-economic models (Dislich et al., 2018; Kirchweger et al., 2020).

While these maps enhance model realism by grounding simulations in actual landscapes, they also introduce several challenges that limit their broader applicability. Maps of real-world landscapes are static objects, often making it difficult to assess how different spatial distributions and patterns within the landscape contribute to the process of interest. Additionally, even if the pattern-process relationships can be disentangled, the unique characteristics of a landscape map taken from a specific location makes the transferability of insight to other locations a difficult matter (Langhammer et al., 2019).

To enhance the generalizability of model insights and disentangle the effects of spatial patterns on agricultural and ecological processes, researchers have developed Agricultural Landscape Generators (ALGs), which generate synthetic maps that mimic real-world landscape mosaics and serve as flexible model inputs (Langhammer et al., 2019). These ALGs can generally be divided into two categories: pattern-based generators and process-based generators. Pattern-based generators are more simplistic and take an algorithmic approach to create geometric properties. In contrast, process-based generators aim at creating landscape patterns by modelling the underlying socio-economic, ecological, and topographic mechanisms and processes that have created current landscape patterns.

Despite their potential, ALGs have remained underutilized due to barriers such as limited accessibility, lack of integration into widely used programming environments, and the inherent tradeoff between generality and realism. The majority of ALGs have not been written in R, the programming language used most by ecologists (Lai et al., 2019). Additionally, ALGs often struggle to balance generality and realism. Many ALGs show high generality and transferability, but they often produce oversimplified landscapes (e.g. Inkoom et al., 2017). This is particularly true for many pattern-based generators. Conversely, there are highly specialized ALGs which can simulate landscapes in great detail, but these cannot be easily generalized or transferred to other locations outside of the region for which they were developed (Hess et al., 2020; Salecker et al., 2019). This mostly applies to process-based generators. While some work has been done towards creating ALGs that maintain high generality while also maintaining a high degree of realism (Etherington et al., 2022; Gaucherel et al., 2014), they still lack an easy pathway for integration in the workflow of the wider modelling community.

To bridge the gap between realism and generalizability, we introduce *ALGR*, a multi-purpose and dynamic landscape generator that combines pattern-based and process-based approaches. It is specifically designed for seamless integration within the R programming environment, ensuring accessibility and flexibility for researchers. Our approach follows a three-step procedure. First, a potential space map is generated, delineating areas available for field placement based on simulated landscape features such as slope, soil, or agricultural constraints. Second, a pattern-based approach is used to place fields within this space. Finally, the landscape is enriched with additional information, including farmer identities and crop distributions. The three-step simulation maintains structural realism, i.e. maps correspond with their real-world counterparts in terms of landscape configuration and geographic properties. This is achieved by mimicking some of the processes of real-world landscape generation (Troost et al., 2023).

Our novel approach allows for a wider use of ALGs in scientific research. To increase its flexibility, our landscape generator includes several methods for generating potential spaces, as well as different algorithms for field placement. *ALGR*, also allows for adding further information to the landscape such as crops, field IDs and farmer information, making it easily integrable with complex ecological models. To increase the accessibility of this landscape generator, we have also encapsulated the code in R package format, which is easily accessible to ecologists interested in using *ALGR* for their own projects.

### The *ALGR* package

*ALGR* generates agricultural landscapes with a three-step approach: first, outlining potential space, second, field placement inside of the potential space, and third, enrichment of spatial information. The functions belonging to the different steps can be seen in Figure 1A, and the example maps resulting from the different functions can be seen in Figure 1B.

**Figure 1A.**
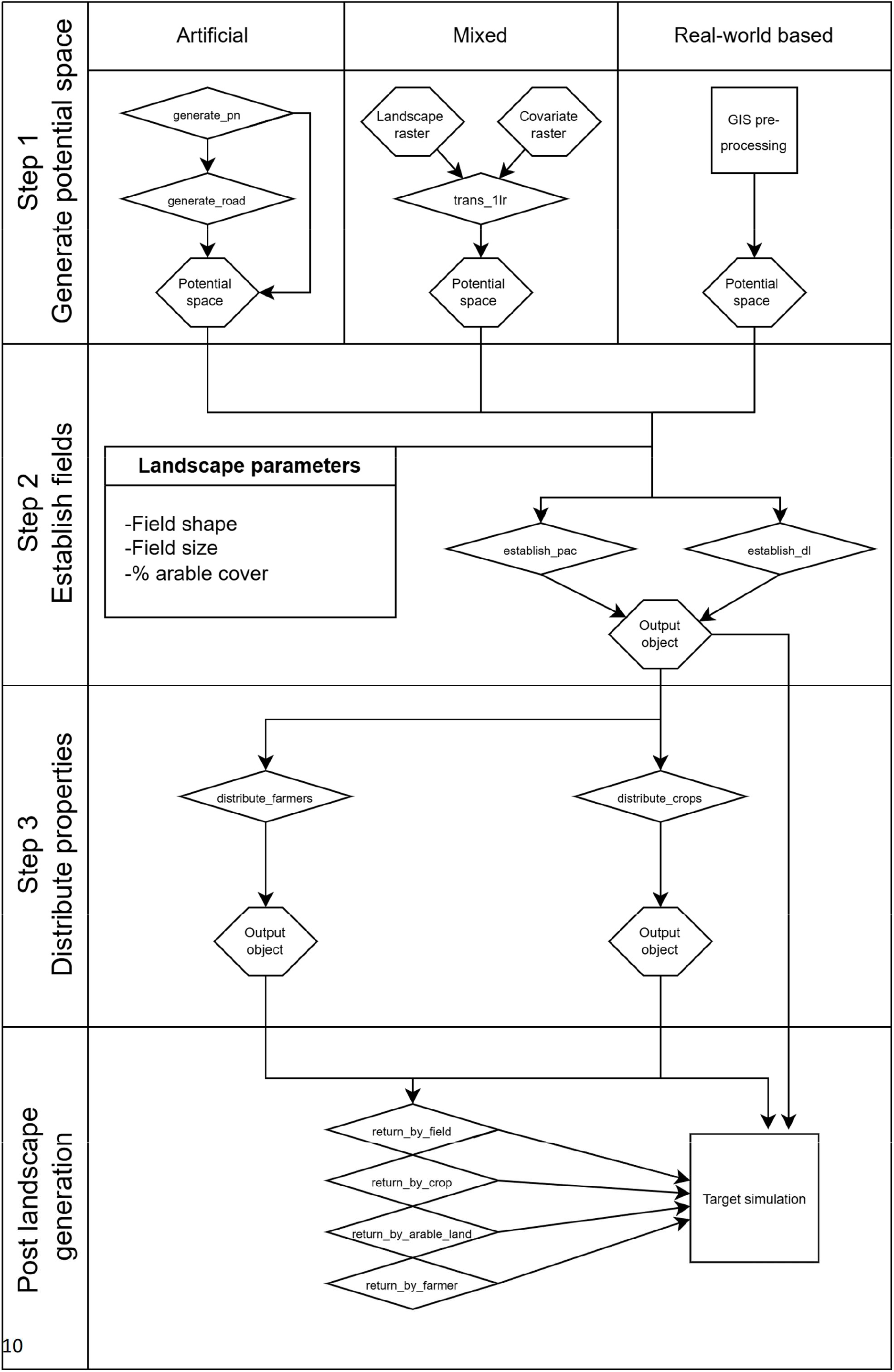
a flowchart illustrating the step-by-step functions of landscape generation: generating potential space, establishing fields inside of the potential space, and distributing properties to the fields. Hexagons: data objects, rhombuses (diamonds): *ALGR* functions, squares: external functions. Additionally, the package includes post-generation functions that enable plotting and extracting various types of raster data from the *ALGR* output object.

**Figure 1B.**
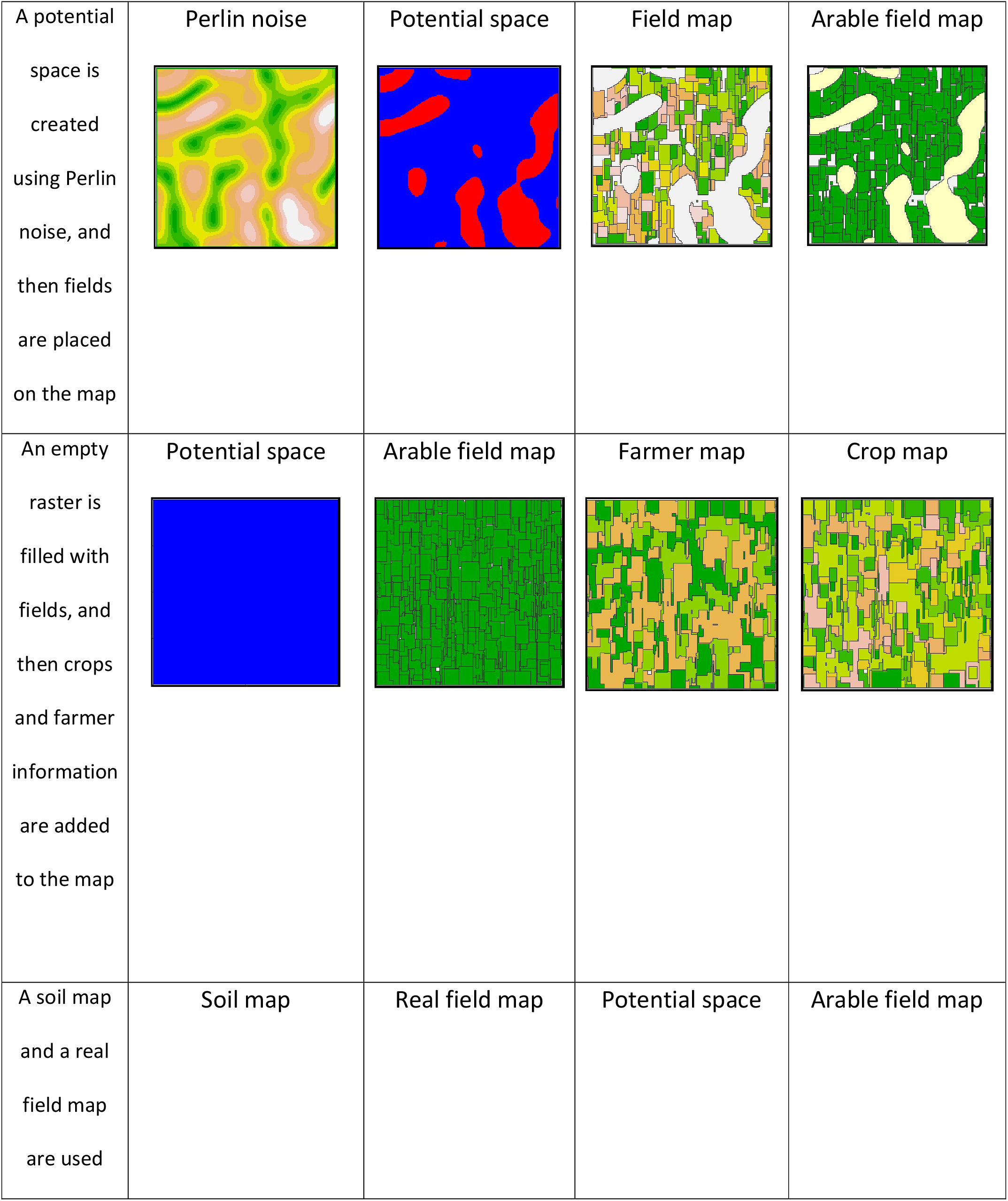
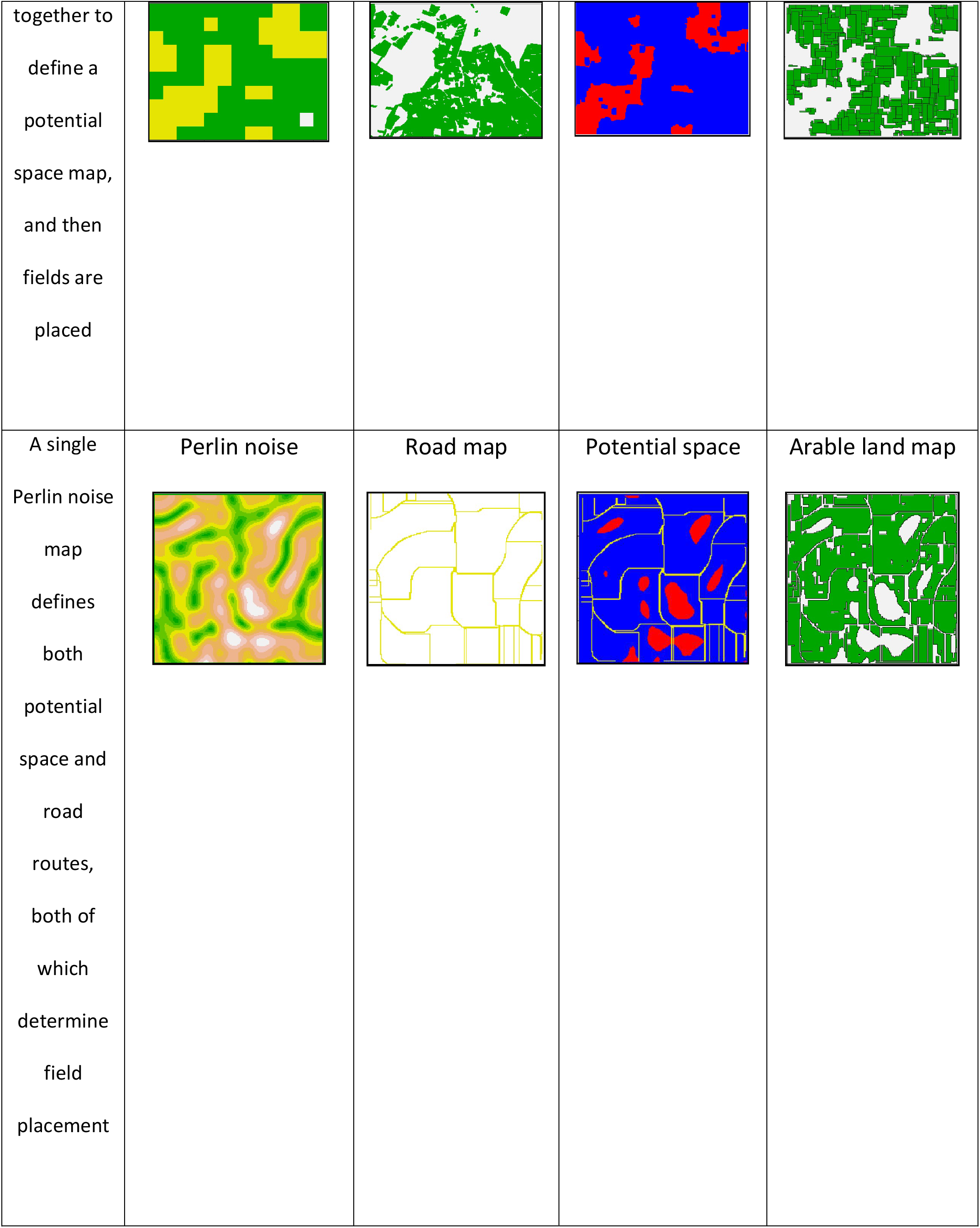
a visual demonstration of the pathways used to generate different types of maps. For each of the potential space maps blue represents available space, while red represents cells that are not part of the potential space. Different output maps are possible, including arable field maps, arable land maps (excluding field borders), crop maps, and famer maps.

### First step: outlining potential space

Three functions are included in the *ALGR* package to outline potential agricultural space. The potential space for arable land can be generated using two built-in methods, Perlin noise or the hybrid world approach. Alternatively, an external raster map with designated arable areas can be imported for use as potential space.

#### Perlin noise approach

The Perlin noise algorithm is an image generator based on the creation of a controlled stochastic effect (Perlin, 1985). It has been widely used for animation (Lagae et al., 2010) and simulation of artificial natural patterns, including terrain (Etherington et al., 2022; Hyttinen et al., 2017). *ALGR* uses a Perlin noise algorithm to stochastically generate a potential space map that simulates terrain-driven agricultural limitation. The flexibility of the Perlin noise algorithm allowed us to also simulate patterns seen in small-scale soil distribution, coarse soil characteristics (referred to as *aglim* in FAO soil categorization), and rough terrain features such as slope and elevation.

The ‘*generate_pn’* function in *ALGR* is a wrapper around the ‘*noise_perlin’* function from the *ambient* package, which is then adapted for specific use. The function takes the raster from the generated by the ‘*noise_perlin’* function and converts it to a map with values ranging from 0-90^°^. This map can then be used in three different ways: 1) it can be categorized into potential space based on a map value threshold specified in the parameters of ‘*generate_pn’*, 2) it can be categorized based on a defined share of potential space cover specified in the parameters of ‘*generate_pn’*, 3) or it can be left as a raw data for further manipulations before being used as input for the second step of the landscape generator.

#### Hybrid world approach

The hybrid world approach models artificial agricultural landscapes within the context of existing terrain. Where a field can be placed (i.e., the potential space) depends on landscape characteristics such as slope, soil, and/or *aglim*. The ‘*trans_1lr*’ function helps determine which areas on the map are likely to be suitable for agriculture. It calculates the probability of each cell in the raster being part of the potential agricultural space, based on two types of input: categorical land cover maps with a single ‘arable landcover’ category and covariate maps. After probabilistically assigning the potential space cells, the function converts neighboring cells by a majority vote to simulate spatial autocorrelation. In other words, the function ensures that areas designated as potential space are clustered together rather than scattered randomly, which results in a potential space area that is larger than just arable land on the original land cover map.

The idea behind this approach is to facilitate the simulation of an arable land configuration in a real-world context. The potential space created using this method contains not only the current location of arable fields, but also possible locations where fields could be placed but are not. Additionally, these maps contain valuable information such as potential space located on suboptimal topography or soil.

#### Additional artificial input: roads and rivers

The package functionality also supports the use of additional raster layers as input, which can further refine the potential space and exclude field placement. This functionality is generally reserved for spatial features such as roads and rivers, allowing them to be included when they are of interest for the output of the generator. We have also provided a tool for artificially generating roads (‘*generate_road’*) and rivers (‘*generate_river’*) using a path of least cost algorithm based on slope maps (either real world slope maps or artificially generated Perlin noise maps).

### Second step: field placement

Second, *ALGR* employs a pattern-based approach to place fields inside the potential space. The package currently includes two different methods for field placements, a place-and-conquer algorithm, and a dead-leaves algorithm. Both algorithms share a set of parameters for field establishment, including: field shape, field size, distribution of size (normal or log-normal), and the percentage of potential space to be covered by fields.

#### Place and conquer algorithm

‘*establish_pac’* uses an algorithm that places each field by segments. The algorithm starts with drawing a field shape value (relation between width and length) and field size value from a distribution defined in the parameters. These values define the length of each segment, and the number of segments to be placed during the establishment of a given field. Each segment is aligned as closely as possible to the previous one, while maximizing both the alignment between both segments and preventing each new segment from being placed on forbidden cells (i.e. outside of the potential space or on another field). This process continues until the field either reaches its pre-set size (all segments have been placed) or there is no more potential space available.

The main advantage of this approach is that it generates very realistic landscapes, where the shape and size of each field is a result of both the parameters and the constraints of the space. The main disadvantage of this method is that it is slightly slower than the dead leaves algorithm for very large landscapes.

#### Dead leaves algorithm

‘*establish_dl*’ is based on a modified dead leaves algorithm (see Galeme & Goussea 2012). In this approach, the potential space is first completely filled using the traditional dead leaves algorithm, including the distribution of randomly drawn rectangular shapes and sizes to be placed inside of the potential space. The modified algorithm then includes a second step where the algorithm starts removing fields beginning from the smallest one, until the percentage of potential space to be covered by fields is reached. The removal of the smallest patches (rather than random patches) is done to avoid the over-fragmentation of the landscape, due to unrealistically many small fields.

The main advantage of this algorithm is its speed. A very large landscape can be generated quickly, but with a cost to realism of the field shape, size, and placement.

### Third step: post-hoc farmer and crop distribution

The *ALGR* package allows for enriching the spatial information by adding crop types and farmer information to the different maps, as well as plotting-tools.

The general structure of the output data from all field establishment functions includes a list of individual fields, with each field containing slots for the following information: field location, field identity, farmer identity, and crop identity. This structure is used to add information to the fields through designated functions but can also be accessed and changed directly.

Farmer identity can be assigned using the function ‘*distribute_farmers’*. The parameters used for this are the mean and standard deviation of the number of fields assigned per farmer and the distribution type (uniform, normal, log normal). The fields are assigned to one farmer at a time until all fields are attributed to farmers. This means that the number of farmers in a landscape is decided by the distribution of fields per farmer and the total number of fields in the landscape, rather than directly parametrized. The fields can be either assigned in a spatially structured way, where nearby fields are given to the same farmer, or in a random way, where the farmer gets random fields from the entire landscape.

Crop allocation to fields is done post-establishment using the ‘*distribute_crops’* function. This function takes a list of crops, and the percentage of arable land they should occupy in the landscape, and then distributes them between the fields. Because only a single crop is assigned to each entire field, larger generated landscapes or those with smaller fields will have better correspondence between the desired cover percentages and the resulting cover percentages in the landscape.

## Example applications

We illustrate the potential of *ALGR* with four examples. The description and results of two examples are included in the main text and the others are in the supplementary information. For each of the examples we included an R notebook in the supplementary information for reproducing the example using the *ALGR* package.

### Example 1: Land use share scenario

Here, we generate landscapes with varying shares of potential space. This potential space is then allocated between arable fields and semi-natural habitat, with both the share and spatial configuration of fields determined by the parameters of the field establishment function. Using landscape metrics, we analyze how changes in the share of potential space and arable fields influence the characteristics of semi-natural habitat patches

In agricultural landscapes, nature conservation policies often focus on conservation by increasing the share of seminatural habitats (Holland et al., 2016). An example is the goal of the EU common agricultural policy to maintain 10% of all agricultural area as seminatural habitat (Pe’er et al., 2020). These seminatural patches can vary in several properties that affect biodiversity and ecosystem services across the landscape, including location within the landscape, size of patches, as well as connectivity and distance between patches of seminatural habitat.

Models dealing with biodiversity and ecosystem services in agricultural landscapes often use real-world maps as an input for simulations (e.g. Holland et al., 2016; Kirchweger et al., 2020). The downside of this approach is that these real-world maps are static objects, making it hard to disentangle the effects of seminatural cover from spatial configuration.

In this example, we show how to use *ALGR* to create a scenario where share of both arable land and seminatural habitat vary within the landscape according to a gradient, along with variation in spatial configuration. Configuration changes arise from two sources: adjustments to field shape and size parameters, and the stochastic effects of different initialization seeds. Starting with a Perlin noise simulation of the topography, we continuously expand the potential space that is available for arable field placement, using ‘*generate_pn’*. This potential space is then divided between arable fields and semi-natural habitat in varying shares using ‘*establish_pac’*.

In this example, the parameters were distributed using a Latin hypercube design, with equal intervals of 1% for the potential space (25-95%), intervals of 1% for the part of the potential space to be covered by arable land (25-95%), as well as 4 field sizes (0.5-2.5 ha and intervals of 0.5 ha), and 4 field shapes which define the ratio between width and length of fields (1-3 and intervals of 0.66).

Finally, we profile the spatial configuration of seminatural habitat patches across the different landscapes using the *landscapemetrics* R package (Hesselbarth et al., 2019). For the spatial configuration profile we use a set of metrices: *mean patch area, largest patch index, fractal dimension, perimeter area ratio, Euclidian nearest neighbor distance*, and *number of patches*.

For the size-related metrics *mean patch area* and *largest patch index* (Figure 2), the largest patches of seminatural habitat were obtained when percentages of potential space and seminatural habitat were the highest (top left section of the plot). For the shape related metrics, fractal dimension and perimeter-area-ratio, the signal was less clear. Fractal dimension of seminatural habitat patches was highest for low percentages of potential space seminatural habitat (bottom right of plot), i.e. when seminatural patches were smallest (see size-related metrics). However, the distribution across the gradient did not show a clear pattern. The perimeter-area ratio of seminatural habitat patches was mostly driven by the percentage of arable cover, with the highest perimeter per area at high arable cover, regardless of the percentage potential space (right side of the plot). The Nearest Neighbor distance showed a pattern similar to the size-related metrics, where the smallest mean nearest neighbor distance between patches of seminatural habitat occurred when percentages of potential space and seminatural cover were highest (top left section of plot). For the number of patches, we found a maximum where the number of seminatural patches was highest when both potential space and arable cover were approximately 90%. Comparable results were achieved for patches of agricultural land (see Figure S1 in section ‘Example 1 – Land share scenario: further results’ of supporting information)

**Figure 2A.**
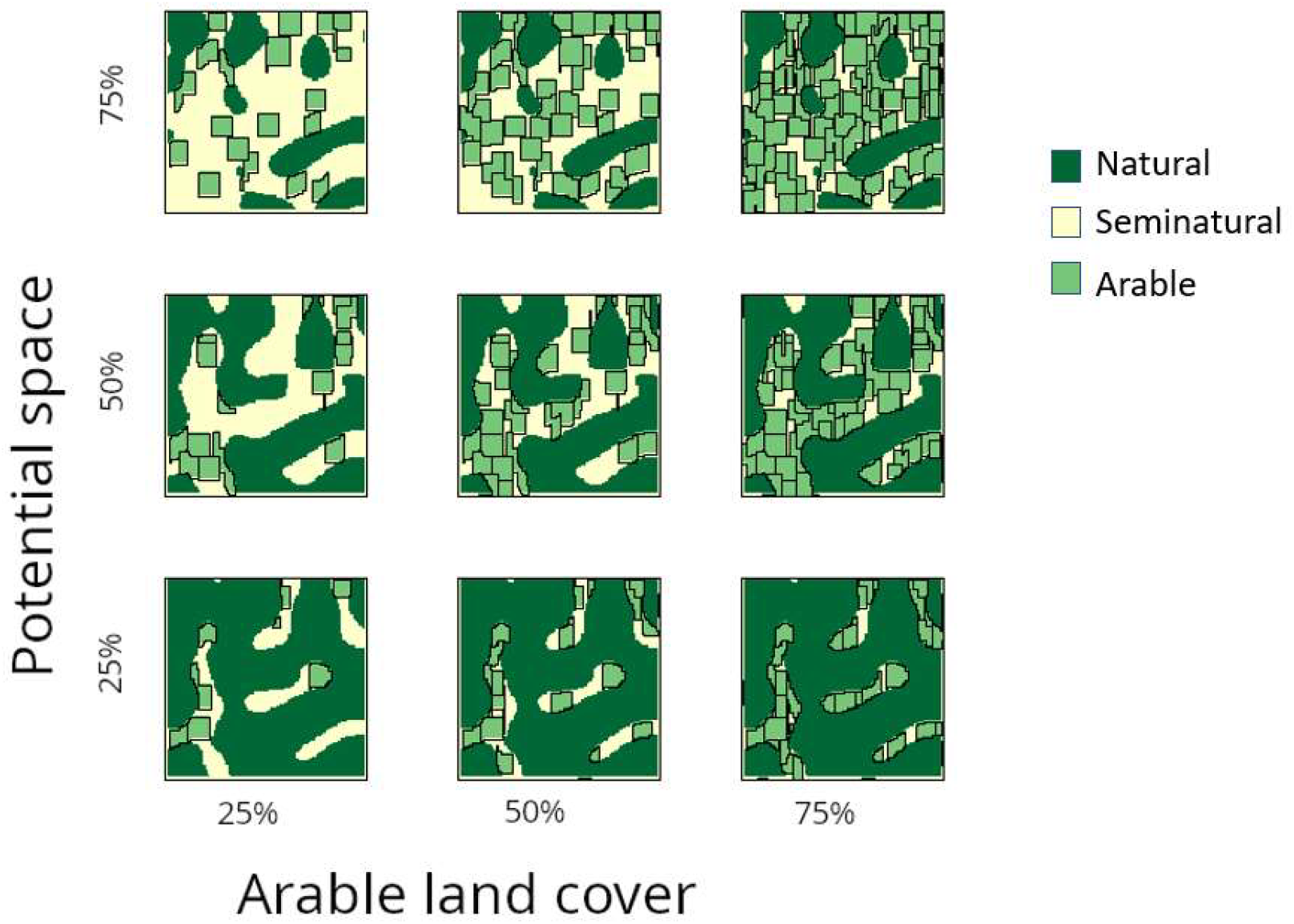
A visualization of the analysis design for example 1, land use share scenario. Landscapes differ along y axis in the percentage of the map available as potential space and along the x axis in the percentage of the potential space occupied by arable land cover.

**Figure 2B.**
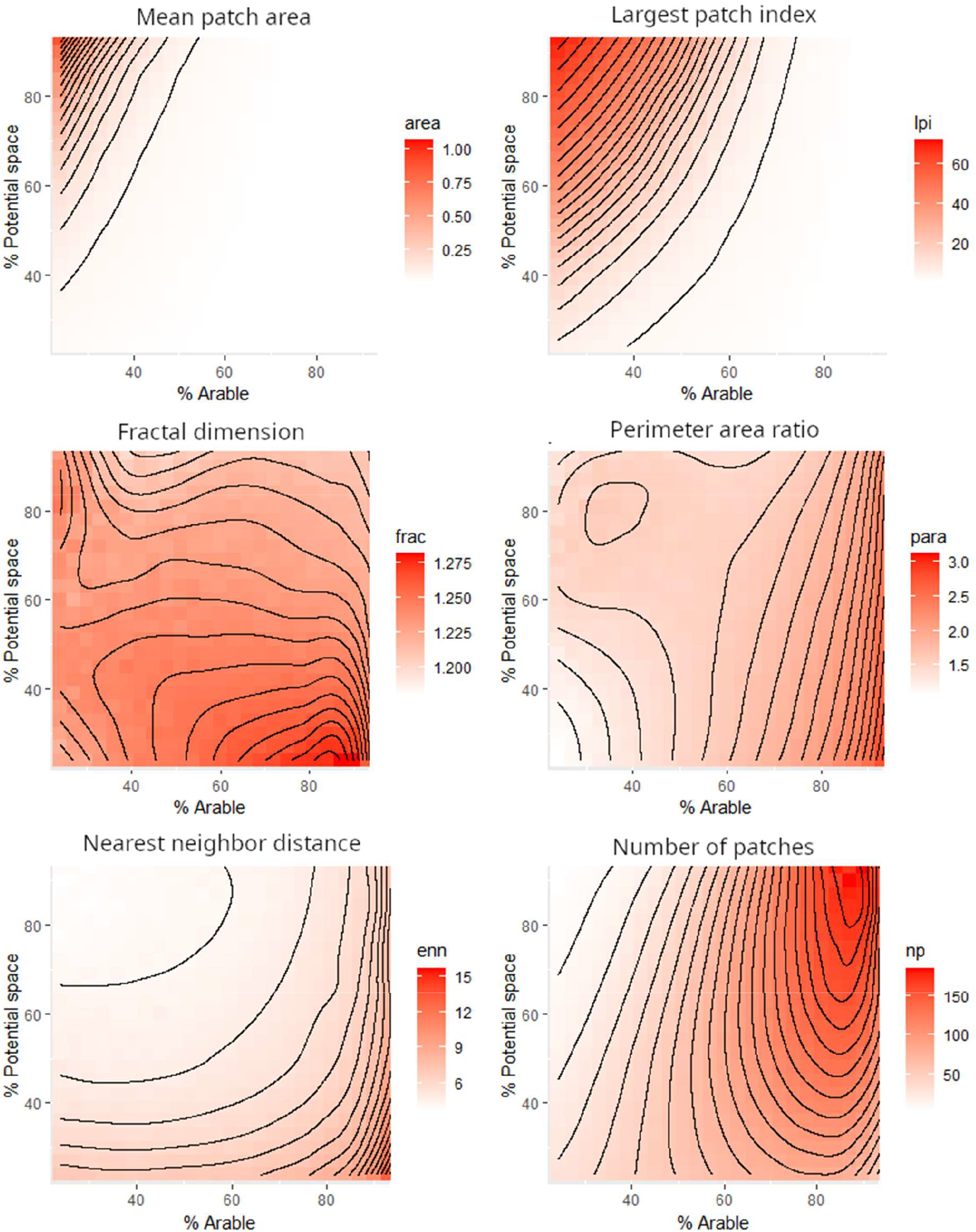
The landscape metrics profile of the seminatural habitat patches across a double gradient where both the potential space and the arable cover vary. For definitions of indices see Hesselbarth et al. (2019).

Most importantly, our results show that that none of the metrics are linear in their response to changes in the share of potential space or arable cover in a landscape. The same change in share of potential space or arable cover will have a different effect on patch structure and configuration, which is often driven by landscape characteristics such as topography. These results are particularly relevant for agricultural policies that promote an increase in seminatural landcover across a range of agricultural landscapes and can be used by modelers to demonstrate differential effects of specific policies.

### Example 2: Pattern reconstruction of agricultural landscapes using a genetic algorithm

Here, we demonstrate how to use a genetic algorithm (Mirjalili, 2019) to fine-tune the parameters of the artificial landscape generator. This process generates maps that share specific characteristics (landscape metrics) with real-world maps.

Landscapes can be characterized by multiple metrics, with each one conveying a certain aspect of the spatial composition and configuration of a landscape (Nowosad & Stepinski, 2019). The spatial features represented by these metrics affect ecological processes, for example fragmentation facilitates metacommunity structure (Hanski & Gilpin, 1991), landscape diversity and simplification can affect ecosystem functions such as pollination or pest control (Tscharntke et al., 2005), and structural connectivity within the landscape can facilitate animal movement (Baguette & Van Dyck, 2007).

Because real landscapes cannot be freely manipulated, it is difficult to disentangle which element of a landscape facilitates a specific ecosystem function of interest. With a simulation approach we can create landscapes with predefined properties and assess their ability to support ecological processes. Here we show how to use a genetic algorithm to parameterize the *ALGR* landscape generator with the goal of recreating the spatial characteristics of a real-world landscape. More specifically, the goal is to keep certain metrics of the landscape fixed, while allowing for variation in other metrics and the overall arrangement. Using the *ALGR* in this way can help modelers generate a diversity of maps while controlling for specific properties of the landscape in each map.

In our example, we used a 2 km × 2 km map as reference. We then used a set of landscape metrics to quantify the optimization goal in our genetic algorithm: the *number of patches* and *patch area* (mean and standard deviation). These two metrics are commonly used in spatial characterization of landscapes. Using the metrics of our reference map as optimization goals, the genetic algorithm searched for an optimal parameterization of *ALGR*. Here, optimal means that *ALGR* generates landscapes that have maximal similarity to the original reference landscape in terms of metric values. Once a parametrization was found (note that several solutions are possible), we can generate any number of maps that are equivalent to the reference map, i.e. similar or even identical in terms of our chosen landscape metrics but different in spatial arrangement (see Figure 3A). We generated 300 maps using the optimal parameterization from the genetic algorithm and compared them to our original reference maps using a list of additional landscape metrics that were not used for the genetic algorithm optimization, including: contagion index (measuring how likely a grid cell of one category will have a neighbor cells of the same category), *largest patch index, fractal dimension, landscape shape index*, and *patch shape index* (see Figure 3B). Our genetic algorithm converged within 30 generations (see Figure S3 in section ‘Example 2 – Pattern reconstruction of agricultural landscapes using a genetic algorithm’ of supporting information).

**Figure 3A.**
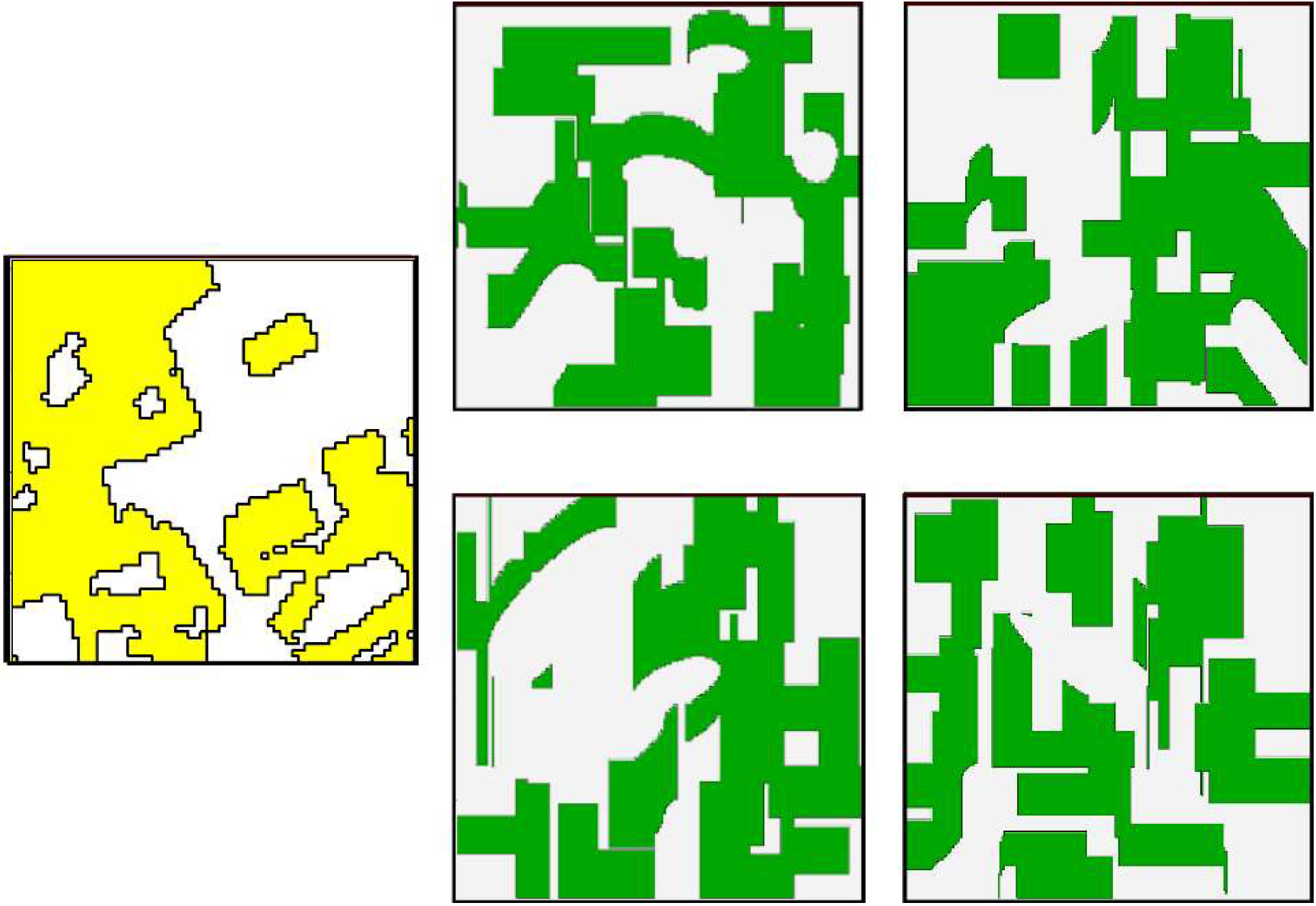
A reconstruction of the landscape pattern of a 2 km × 2 km landscape using the metrics *number of patches* and *patch area*. Yellow: reference landscape used as optimization goal for our genetic algorithm. Green: four realizations obtained using the optimized *ALGR* parameters.

**Figure 3B.**
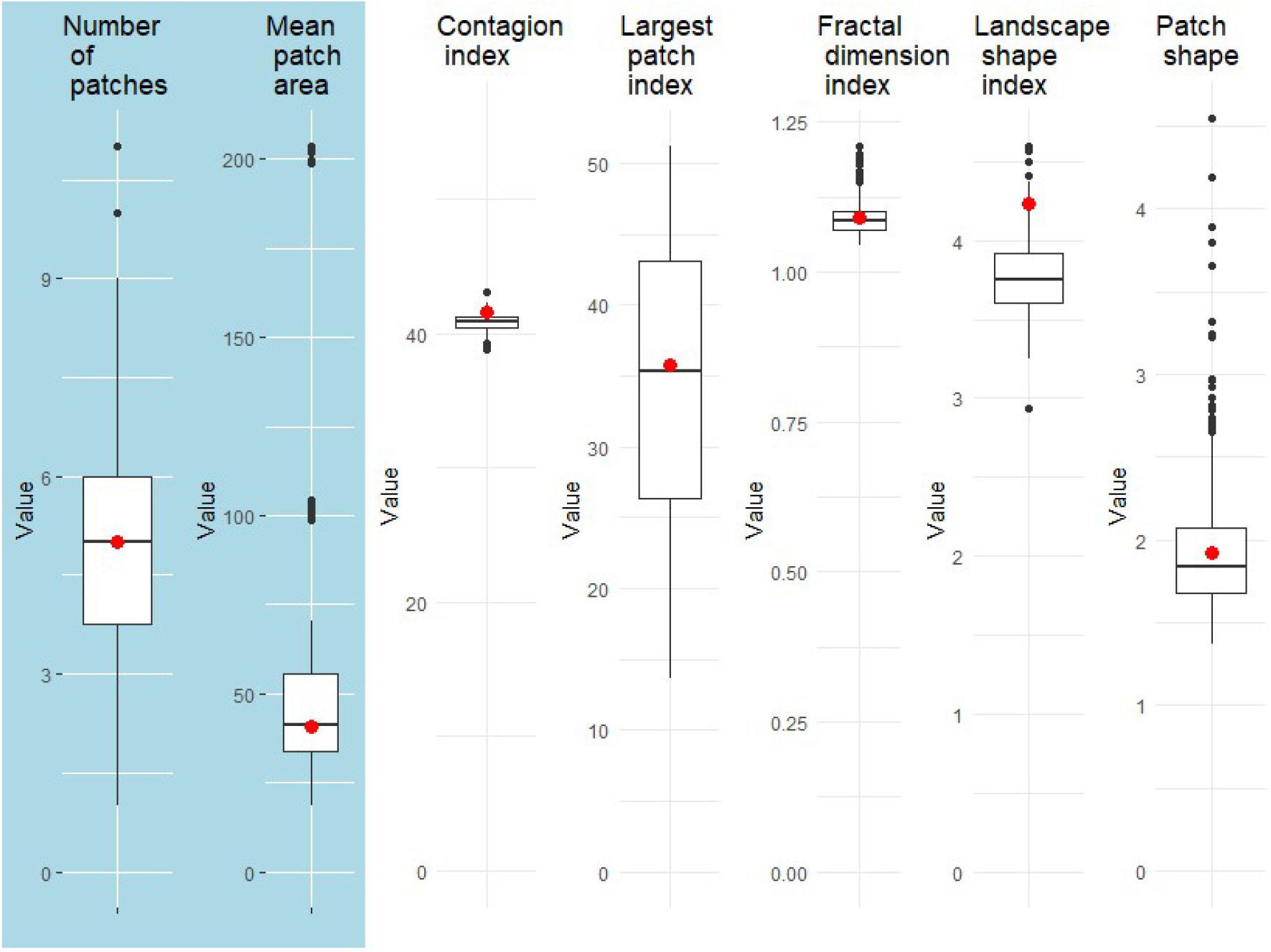
A comparison of different landscape metrics between our 300 generated maps (boxplots) and the reference landscape (red dots). Blue background: metrices that were used as optimization goals in our genetic algorithm, including number of patches and mean patch area (ha), and white background: additional metrices that were not part of the optimization.

### Example 3: Crop portfolio to landscape scenario

This example demonstrates how non-spatial crop share portfolios are translated into spatially explicit, artificially generated landscapes. We then use simple landscape metrics to demonstrate how these different realizations might affect a spatially explicit ecological process (see section ‘Example 3: Land use allocation problem’ in supporting information)

### Example 4: Real world maps as potential space

This example includes the use of real-world maps for the potential space and demonstrates how to generate fields on top of the potential space (see section ‘Example4: using real-world maps as basis for landscape simulation’ in supporting information)

## Conclusion and outlook

*ALGR* is the first agricultural landscape generator readily available within the framework of R that allows for systematic and reproducible landscape generation. It can be used to simulate maps needed for agricultural and ecological models that require landcover maps as input, allowing for generality of use across different contexts and regions while maintaining structural realism. While other ALGs have been previously published (Langhammer et al., 2019), *ALGR* has the advantage of being available in open-source format and well-integrated in the R workflow, the most prevalent programming environment used by ecologists.

The *ALGR* package is specifically designed to facilitate the different needs of ecologists and modelers interested in using an ALG in their workflow. Besides the multiple functionalities presented in this paper, the package is designed to always output an object that stores both field information as well as farmer information, therefor allowing the maps to be easily integrated with more sophisticated ecological and socio-economic models, as well as be further manipulated using simple R code.

Though *ALGR* is general in its application, it was developed with European agricultural landscapes in mind. In these regions, the configuration of agricultural landscapes is often determined by the availability of ‘suitable’ space for arable land. Thus, in its current version, other factors are not included which are of higher importance in other regions, such as proximity to water bodies (e.g., North Africa) or roads (e.g., Indonesia, Brazil).

To facilitate the use of the package, we have included a notebook to reproduce each one of the examples presented in this paper. The help sections also include working examples of all functions. Finally, we have also included a vignette that provides a full description of each one of the functions included in the package which is also hosted on the same github repository.

## Data availability statement

The code for *ALGR* was written in R (R Core Team, 2024) and is openly available to download from the following GitHub depository (https://github.com/pogoyoly/ALGR) free to be used and further expanded. The datasets used for all examples of the model application presented here are all open source and freely available.

## Author contributions

Eyal Goldstein, Antonia Deutscher and Eamon O’Keeffe developed the *ALGR* R package. Kerstin Wiegand was the principle investigator. Eyal Goldstein was the lead author of the manuscript, and all authors contributed critically to the drafts and gave final approval for publication.

