## supporting information for "*ALGR*: A multi-purpose agricultural landscape generator in R"

**Supplementary information on paper, ALGR: A multi-purpose agricultural landscape generator in R**

**Example 1 – Land share scenario: further results**

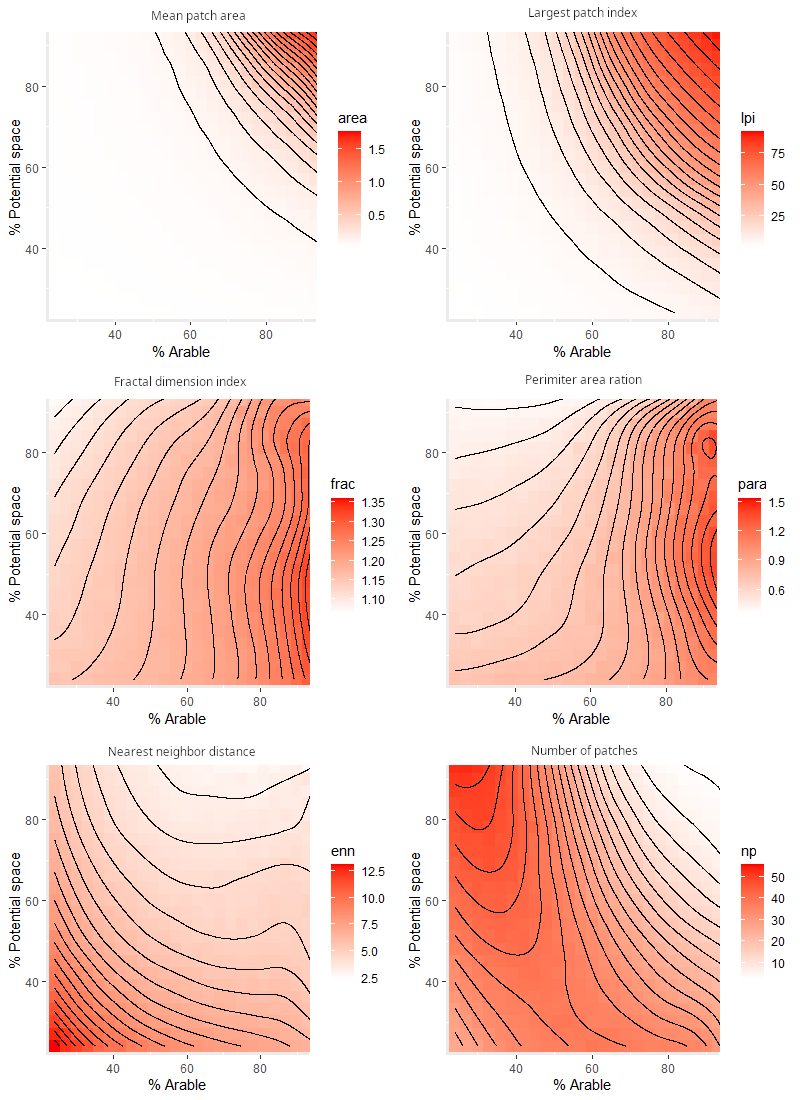

Figure S1: Landscape metrics profiles of the arable land patches across a gradient where both the potential space (% Potential space) and the arable cover (% Arable) vary between simulations.

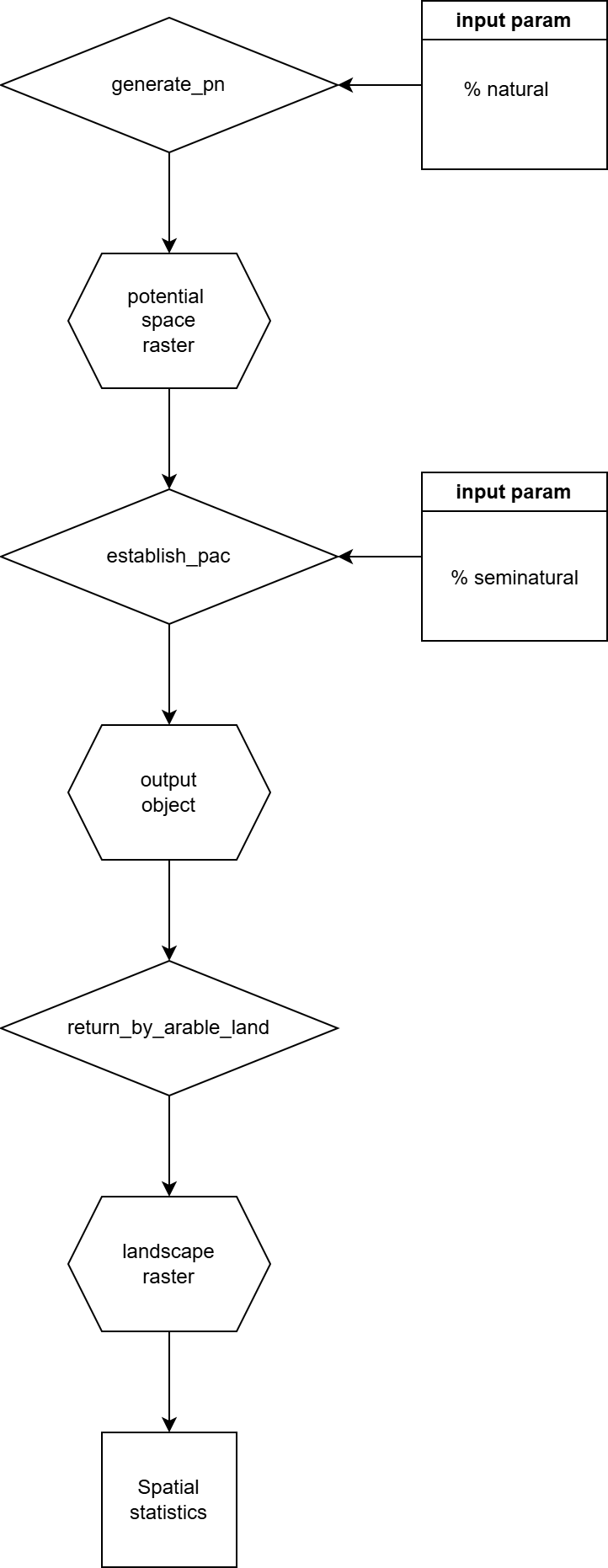

Figure S2: A flow chart of the different functions and objects used during example 1

**Example 2 – Pattern reconstruction of agricultural landscapes using a genetic algorithm** **: further results**

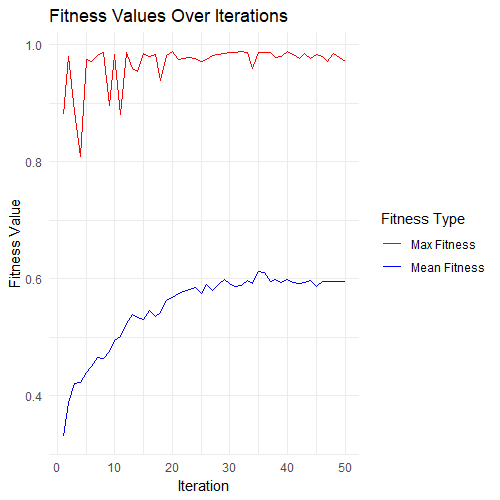

Figure S3: evaluation of the fitness function for our optimization of parameters in example 3A-B

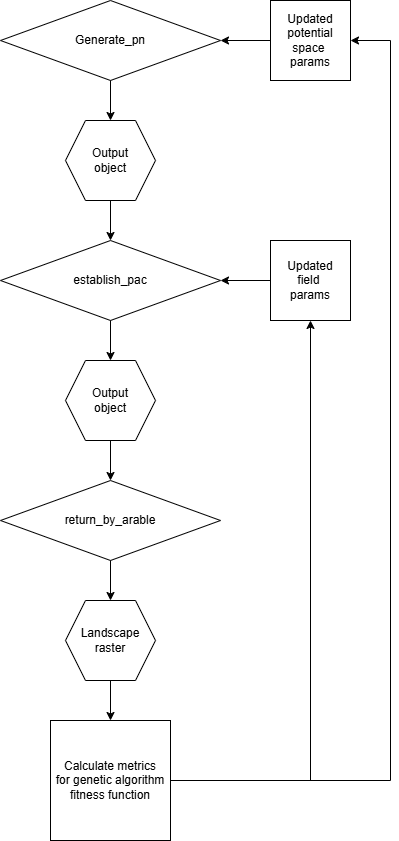

Figure S4: A flow chart of the different functions and objects used during example 2

**Example 3: Land use allocation problem**

Land use allocation according to different criteria is a known challenge in sustainable development (Kaim et al., 2018) and a general question for agricultural economists. Land use portfolio optimization, in other words finding the share of different land use categories which together best provision for different goals or purposes, is a common tool in land use multicriteria decision making. One major drawback for this method is the lack of consideration for space during optimization, as well as lack of spatial representation of its results. In ecology, in particular spatial representation is important because many ecological processes are spatially dependent (Hanski, 1998; Tscharntke et al., 2005), for example wild pollinators can use different land cover categories for feeding and nesting, and their ability to provide pollination services is influenced by the landscape structure (Tscharntke et al., 2012).

In this example, we show how to transform non-spatial land-use portfolios to spatially-explicit maps using ALGR. The portfolios that we used (Cong et al., 2014) were exclusively optimized for profit under different risk values. Risk is defined as the maximum accepted variance of profit return (see Figure 5 for portfolio composition). These portfolios were the so-called ‘statics’ portfolios in Cong et al. (2014) (see Figure 3 in Cong et al. 2014). The values were extracted using a graph digitization software (<https://automeris.io/>). We then generated landscapes using the ‘*establish_pac’* function, completely filling up the landscape with fields (100% potential space and 100% field placement). We then used the ‘*distribute_crops*’ function to allocate the crop types according to their share in the portfolio.

To analyze the implications of the different portfolios for pollination ecosystem services, we focused on the distance between feeding resource cells (rapeseed cells) and nesting resource cells (grassland cells). While rapeseed requires (wild or domesticated) bees for pollination, it does not provide good nesting resources for wild bees (citation). Grasslands, on the other hand, can provide a good nesting resource, but the rapeseed fields must be within a certain, species-dependent distance for bees to provide adequate pollination. To estimate levels of pollination, we calculated the mean distance to the nearest grassland cell for all rapeseed cells. For each portfolio, 5000 landscape realizations were generated across a gradient of varying field sizes. For each landscape, we calculated for each rapeseed cell the distance to the nearest grassland cell. We plotted the results as the mean across the mean-nearest-distance of all landscapes and estimated 95% confidence intervals using the lower 2.5% and upper 97.5% quantiles.

Our results show significant differences between portfolios in terms of pollination, as estimated for each landscape via the mean nearest distance to grassland cells for each rapeseed cell (See figure S5C). There was a clear trade-off between portfolio risk and pollination. Portfolios calculated for high-risk high-return (3/4 Var) showed lower mean minimal distance to nearest grassland cells. This was mostly because grassland was a low-risk low-return land use category in the Cong et al. 2014 static model, which lead to a decrease in grassland as the risk level was increased. For the high-risk portfolios as well as for the observed portfolio, field size had a strong effect on the increase in mean minimal distance, highlighting the importance of multiple spatial consideration for a full assessment of land use composition effects on spatially explicit ecological processes. Our simulation results also demonstrate new levels of tradeoff beyond just risk and return, which can be calculated using landscape simulation methods.

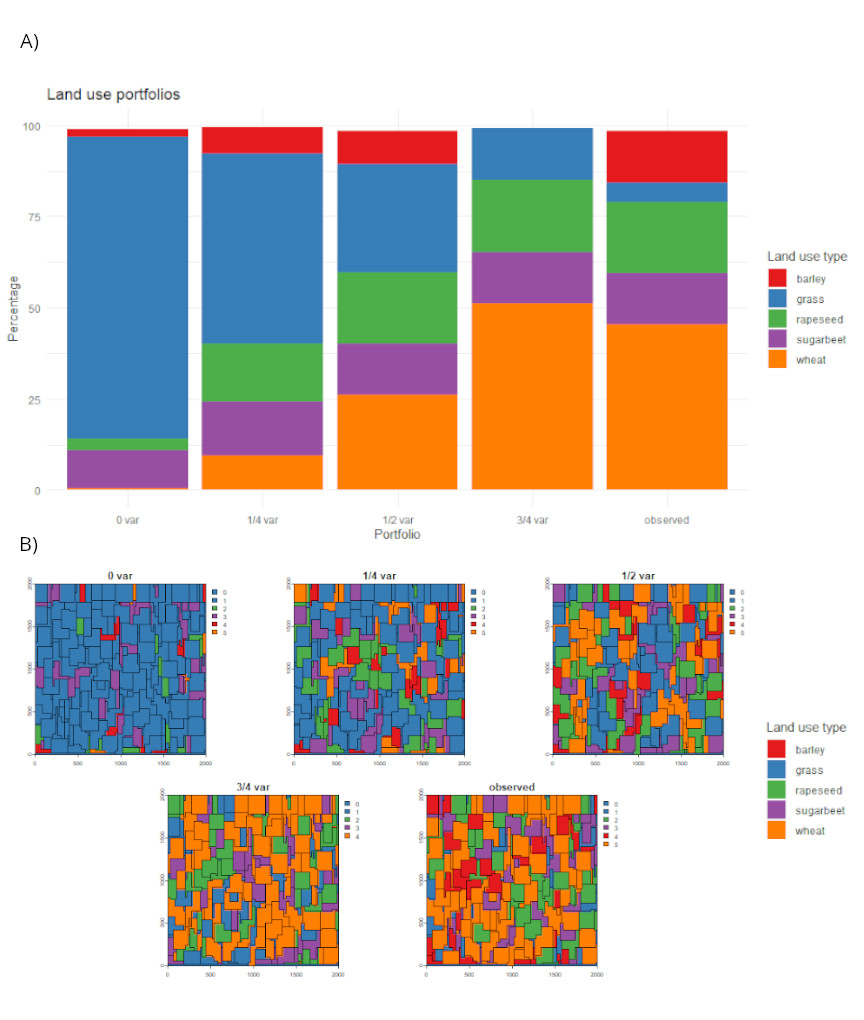

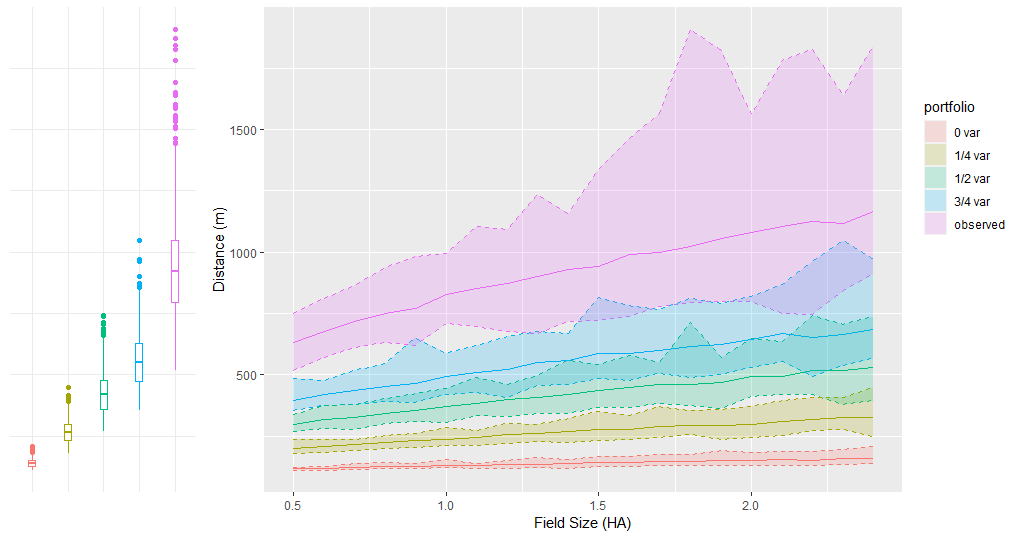

Figure S5: A) bar plot of each portfolio composition used in this example, B) one example of spatially explicit representation for each portfolio generated using the ALGR package and a field size of 0.5-3 ha, C) Mean distance between each rapeseed cell and the nearest grassland cell for the five portfolios (see legend). The boxplots lower hinge representing the 25^th^ percentile and the upper hinge representing the 74^th^ percentile, and the whiskers representing extend to 1.5 of the interquartile range. The continuous function (right) represents the mean distance (continues line) and the lower and upper bound of mean distance using the 1^st^ and 99^th^ percentile. The mean distance generally increases as the field size increases.

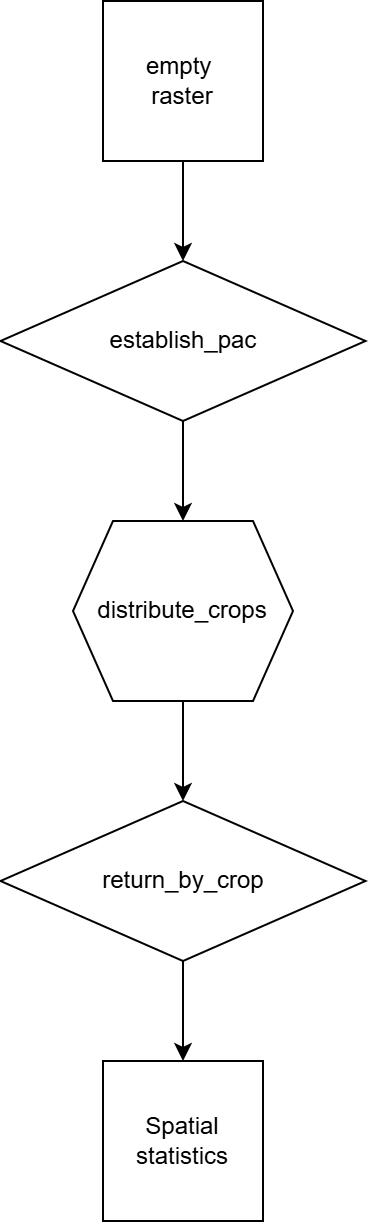

Figure S6: A flow chart of the different functions and objects used during example 3

**Example4: using real-world maps as basis for landscape simulation**

In the following example, we demonstrate how real world landcover maps can be used along with ALGR to simulate landscapes. For this example, we took a 12 km × 12 km section of the Copernicus EU landcover map from southern Germany that shows the dominant landcover category for each 100 m × 100 m raster cell. The maps that we used had three different landcover categories: forest, grassland, and arable land. In our example we considered all cells marked as arable land as potential space. When then took 2 km × 2 km sections of the map and downscaled them to a spatial resolution of 10 m × 10 m. Finally, we used the map sections as input for the *‘establish_pac’* function, which places fields across the potential space.

Our results demonstrate the possibility of using an external raster as input for ALGR (As shown in Figure 7A-C). This option can be particularly useful when there are certain unmovable categories (e.g. forest and grassland), but the cover and configuration of arable fields is changeable. Each one of the examples in S7C can be change in configuration by changing field sizes, shape, share of arable landcover, and change in initialization seed.

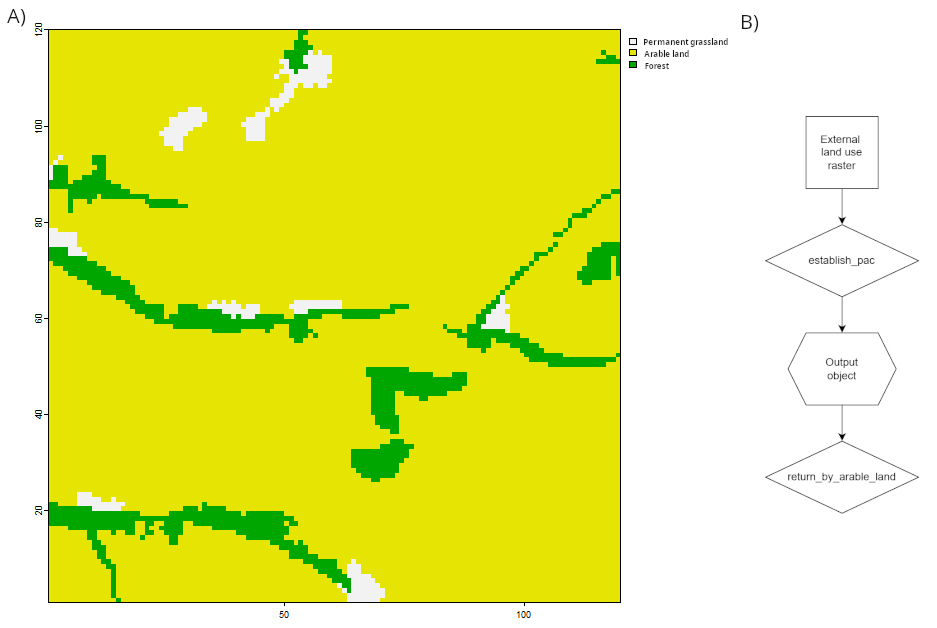

S7: A) A 12km x 12km map of dominant land use categories taken from the Copernicus landcover dataset. B) The flowchart for using an external map as input for ALGR. C) example of field placements on the ‘arable’ land use category of the Copernicus input map.

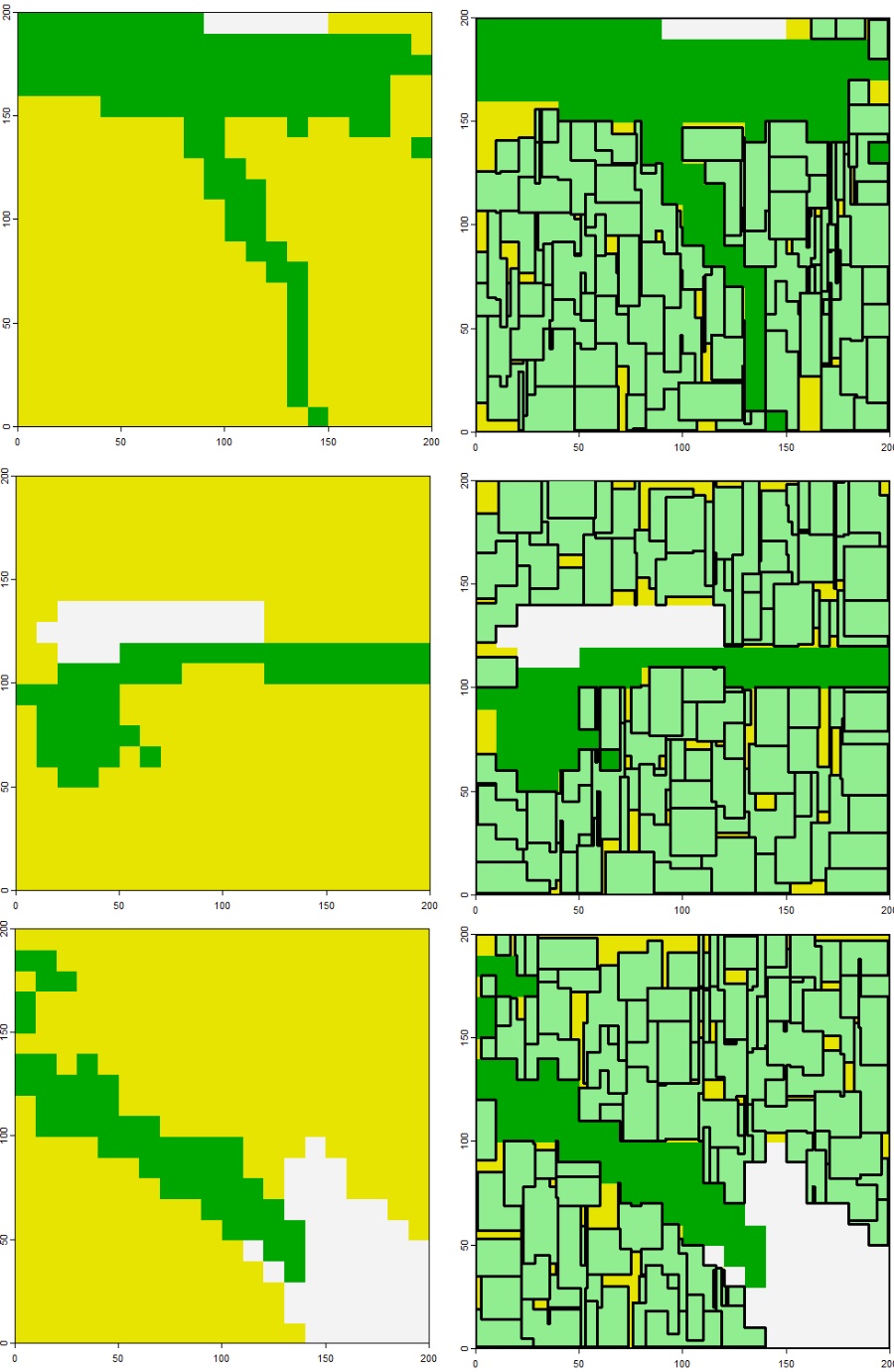

Figure S7: C) example of field placements on the ‘arable’ land use category of the Copernicus input map, and the ALGR function ‘*establish_pac’*
